# Transcranial Focused Ultrasound Generates Skull-Conducted Shear Waves: Computational Model and Implications for Neuromodulation

**DOI:** 10.1101/2020.04.16.045237

**Authors:** Hossein Salahshoor, Mikhail G. Shapiro, Michael Ortiz

**Affiliations:** Division of Engineering and Applied Science, California Institute of Technology, Pasadena, CA 91125, USA; Division of Chemistry and Chemical Engineering, California Institute of Technology, Pasadena, CA 91125, USA

## Abstract

Focused ultrasound (FUS) is an established technique for non-invasive surgery and has recently attracted considerable attention as a potential method for non-invasive neuromodulation. While the pressure waves generated by FUS in this context have been extensively studied, the accompanying shear waves are often neglected due to the relatively high shear compliance of soft tissues. However, in bony structures such as the skull, acoustic pressure can also induce significant shear waves that could propagate outside the ultrasound focus. Here, we investigate wave propagation in the human cranium by means of a finite-element model that accounts for the anatomy, elasticity and viscoelasticity of the skull and brain. We show that, when a region on the frontal lobe is subjected to FUS, the skull acts as a wave guide for shear waves, resulting in their propagation to off-target structures such as the cochlea. This effect helps explain the off-target auditory responses observed during neuromodulation experiments and informs the development of mitigation and sham control strategies.

---

Focused ultrasound (FUS) is an established therapeutic modality taking advantage of the ability of sound waves to deliver energy to anatomically precise regions of organs such as the human brain [1–3]. Recently, low-intensity transcranial FUS has elicited growing interest as a tool for neuromodulation, owing to its concurrent benefits of relative safety, non-invasiveness and millimeter-scale precision [11–22]. However, the biophysical mechanisms underlying neuromodulation are not well understood, and recent studies have documented off-target auditory responses to FUS neuromodulation in rodents [23, 24] and humans [25]. To better understand these phenomena at both the tissue and cellular levels, computational models can play a useful role [26, 27].

Modeling ultrasound wave propagation in the brain requires realistic models that accurately represent anatomical detail and the mechanical response of the tissues. In recent years, detailed computational models have been successfully constructed from magnetic resonance (MR) images [28–31]. The constitutive modeling of soft biological tissues has also received considerable attention [32, 37]. Due to the complexity of the mechanical response of the tissues, the material parameters reported in the literature differ by orders of magnitude [35, 38]. These uncertainties notwithstanding, the large contrast between the bulk and shear modulus is generally understood to result in widely disparate longitudinal and transverse wave speeds, with the former in the range of 1000 to 1500 m/s and the latter at most 10 m/s [39, 41]. Moreover, shear waves are strongly attenuated in soft biological tissues [39, 41–44]. This shear compliance and strong shear wave attenuation properties often allow soft tissues to be modeled as acoustic media [26]. By contrast, this assumption fails in the presence of hard structures such as bone, which can sustain shear waves of amplitude comparable to pressure waves and can act as waveguides by virtue of their extreme impedance contrast to soft tissues.

In the present work, we investigate wave propagation in the human cranium by means of a finite-element model that accounts for the anatomy, elasticity and viscoelasticity of the skull and brain (**Fig. 1**). We employ a high-resolution solid model from the SCI Head Model project [45] constructed from MR images obtained from a 23-year old healthy female subject. The model comprises the scalp, skull, cerebrospinal fluid (CSF), grey matter (GM), white matter (WM), eyes, ears and sinuses (**Fig. 1**).

**Fig. 1.**
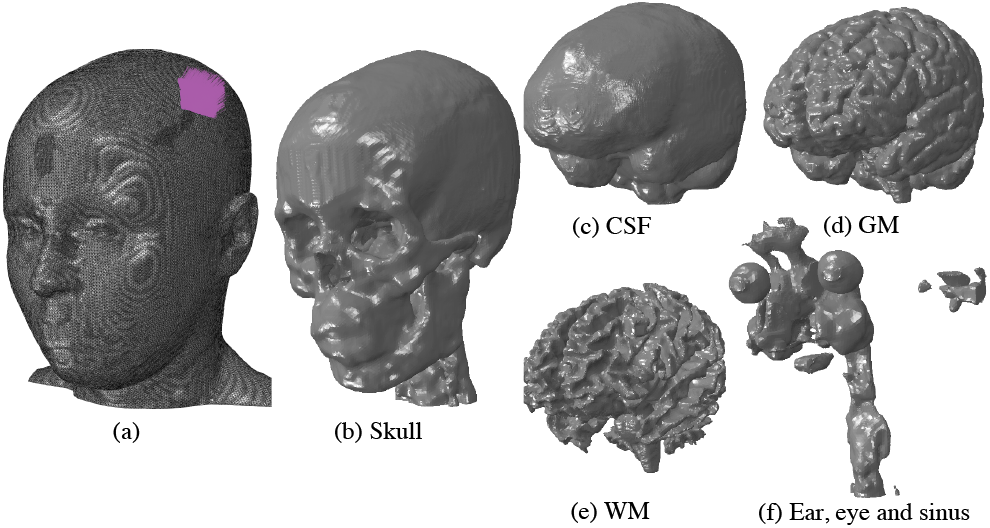
High-resolution solid mechanical model of the human cranium from the SCI Head Model project [45]. (a) Total model with 8,512,657 nodes and 48,458,912 million linear tetrahedral elements. The region on the scalp subjected to ultrasound pressure as traction boundary is shown by purple arrows. Inner parts of the model include the (b) skull, (c) cerebrospinal fluid (CSF), (d) grey matter (GM), (e) white matter (WM) and (f) combined ear, eye and sinus.

On this domain, we solve the initial boundary-value problem of small-strain viscoelasticity. Finite elasticity models have been investigated [36] and found to be indistinguishable from small-strain Hookean models under the low-intensity FUS conditions of interest here. The material parameters used in calculations for the various tissue types in the model are taken from literature [46, 47] and collected in **Table 1**. In this table, κ is the bulk modulus, G is the shear modulus and *ρ* is the mass density. The six-order of magnitude contrast in the shear moduli of bone and soft cerebrospinal tissue is remarkable, as is the similar contrast between the bulk and shear moduli in the soft tissue. The discrepancy between bulk and shear moduli in the soft tissue is often taken as a basis for neglecting shear waves and accounting for pressure or sound waves only [26]. However, when the entire skull/brain system is considered in its entirety, shear stiffness and impedance mismatch strongly influence wave patterns. The viscoelastic properties of the soft tissues are modelled by means of the standard linear solid model with exponential relaxation function:

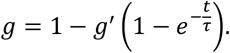

**Table 1:** Elastic and viscoelastic properties of different tissues of the head where κ is the bulk modulus, G is the shear modulus and ρ is the mass density.

| | $\kappa(\text{Pa})$ | $G(\text{Pa})$ | $\rho (\text{N/m}^3)$ | $g'$ | $\tau(\text{s})$ |
| --- | --- | --- | --- | --- | --- |
| Skull | 4.76e+9 | 3.28e+9 | 1721 | --- | --- |
| Scalp | 3.36e+9 | 6.7e+5 | 1100 | 0.6 | 3e-5 |
| GM | 1.2e+9 | 1.2e+3 | 1060 | 0.8 | 80 |
| WM | 1.5e+9 | 1.5e+3 | 1060 | 0.8 | 80 |
| CSF | 1.33e+9 | 20 | 1040 | --- | --- |
| Ear/Sinus | 8.33e+5 | 3.85e+5 | 1000 | --- | --- |
| Eye | 1.13e+7 | 2.28e+3 | 1078 | --- | --- |

The values of the relaxation parameters are taken from [46, 47] and shown in **Table 1**. The CSF is approximated as an elastic medium with an exceedingly small shear modulus but capable of transmitting pressure waves.

The skull and brain geometry is discretized into a finite-element model comprising 48.4 million three-dimensional tetrahedral elements and 8.5 million nodes (**Fig. 1)**. We subject a region in the frontal lobe of the scalp, shown in purple in Fig. 1, to sinusoidal pressure with a peak amplitude of 0.6 MPa and frequency of 200 kHz. Acoustic focusing is modeled by imposing the applied pressure with a phase offset in the radial direction. The governing equations are integrated in time by means of the explicit Newmark algorithm, as implemented in the commercial code Abaqus/Explicit (Dassault Systemes Simulia, France).

Representative computed transient pressure and shear wave patterns (**Fig. 2** and **Fig. 3**, respectively) depict contours of pressure and Mises stress at four different times after the onset of FUS. The results are shown in a frontal cross section of the head through the center of the area of application of FUS. These results show that the pressure wave is transmitted through the skull to the soft tissue and propagates through the brain. At the same time, shear stresses inside the brain have very low amplitudes, on the order of a few Pa, and propagate significantly more slowly. As expected for low-intensity FUS, the computed pressures are below the values reported for injury thresholds for pressure and shear stress [48–52].

**Fig. 2.**
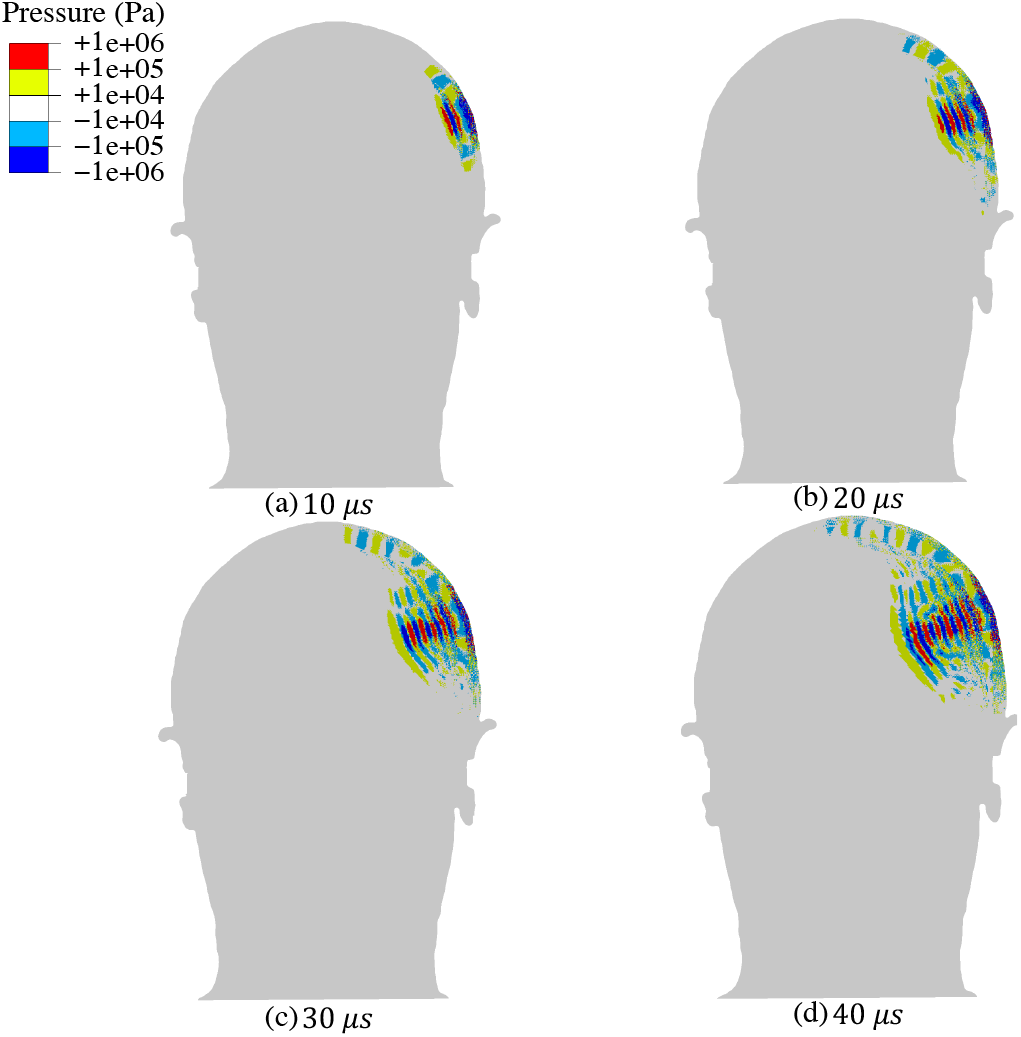
Transient pressure wave propagation due to the application of continuous sinusoidal ultrasound with an amplitude of 0.6 MPa and frequency of 200 kHz to a region of the frontal lobe of the human head. The snapshots correspond to the pressure distribution in a cross-section of the head including the ultrasound focus at: (a) 10 *μs;* (b) 20 *μs;* (c) 30 *μs* and (d) 40 *μs*.

In contrast to the transmission of pressure waves to the brain, our model shows that shear waves are guided by the skull (**Fig. 3**), suggesting that that the skull acts as a shear waveguide. This mechanism results in the conduction of shear waves to locations well outside the ultrasound beam path. By 30 μs after the start of FUS application, the shear waves reach the cochlea, applying stress on the order of 100 Pa to this auditory organ. The rapid transmission of shear waves to the cochlea may help explain the off-target auditory responses recorded during neuromodulation experiments [23, 24, 25].

**Fig. 3.**
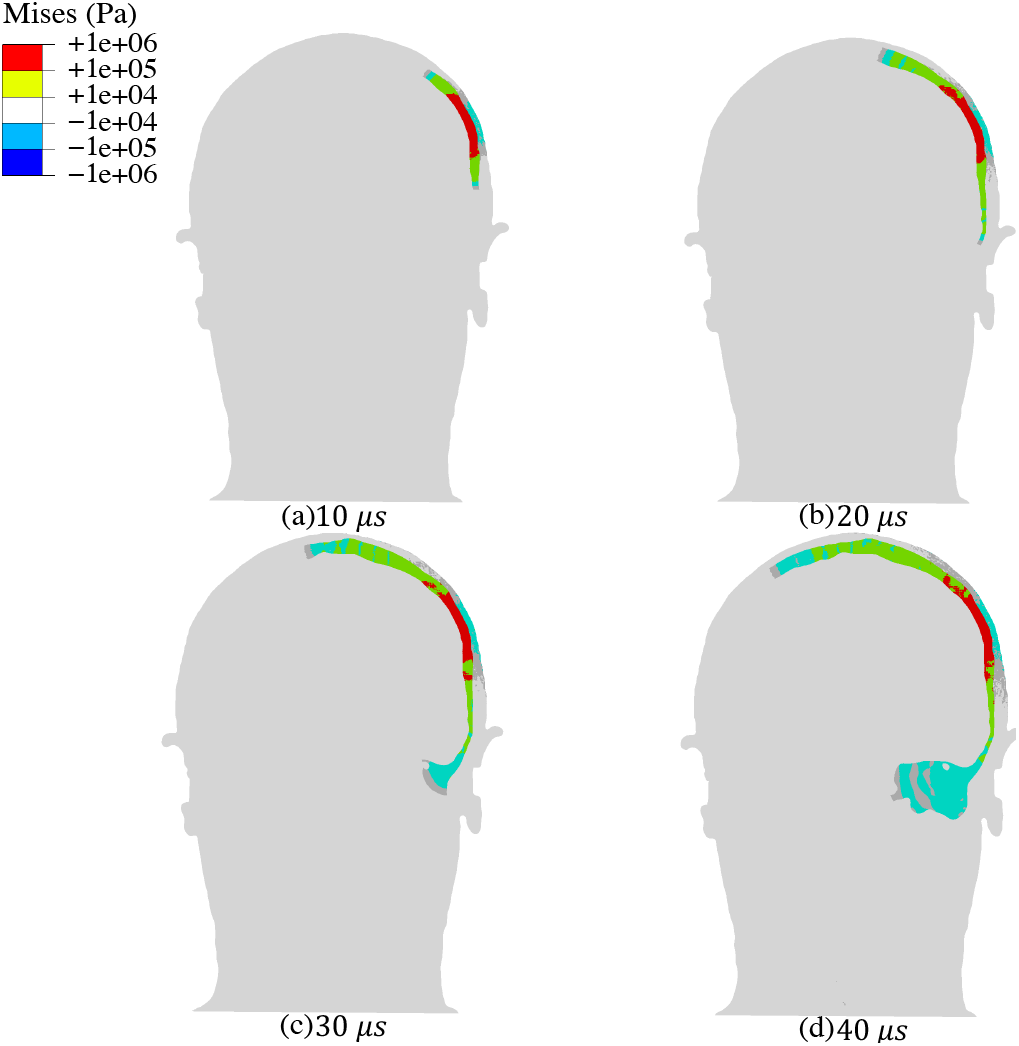
Transient shear wave propagation due to the application of continuous sinusoidal ultrasound with an amplitude of 0.6 MPa and frequency of 200 kHz to a region of the frontal lobe of human head. Bone conduction of shear waves through the skull and towards inner ear is observed. The snapshots correspond to the von Mises stress distribution in a cross section of the head including the ultrasound focus at: (a) 10 *μs;* (b) 20 *μs;* (c) 30 *μs* and (d) 40 *μs*.

Several recent studies have argued that FUS waveforms can be designed to mitigate auditory side-effects. For example, gradual ramping of the applied wave amplitude is proposed to reduce the generation of audible frequencies in the ears [54–55]. To investigate this possibility while assessing the utility of our computational model in pulse waveform design, we subjected the model to two distinct FUS profiles. In the sharp profile, we applied a 200 kHz waveform with an immediate amplitude of 0.6 MPa for a stimulation time of 0.5 ms (**Fig. 4a**). In a ramped profile, we gradually increase the amplitude, reaching 0.6 MPa pressure over 0.1 ms (**Fig. 4b**). The total stimulation time in the second waveform was extended such that the total pulse energy is identical in both cases. With recourse to the aforementioned methods, we computed the displacements in a zone of the inner ear for both cases (**Fig. 4c**). The envelope of a family of curves, where each curve corresponds to the displacement magnitude of a node in the inner as a function of time, is plotted for both scenarios in **Fig. 4d**. The displacements resulting from sharply applied FUS reached a magnitude of 1 μm, coinciding with the displacement ranges in the Stapes and Basilar membrane needed for bone conduction hearing [56]. Meanwhile, the ramped pulse produces a maximal displacement of 0.2 μm. This difference in displacements is consistent with the general response of an elastic system to step and ramp functions, pertinent to the theory of oscillations [57]. While we cannot conclude that the 5-fold reduced magnitude of displacement in the ramped pulse is low enough to eliminate auditory effects, our finding provides support for ramping in general as an approach to reducing them. Due to computational limitations, the ramping time used here was shorter than those used in some neuromodulation studies [54–55]. Such more gradual ramps may be expected to further reduce the maximal shear-generated displacement. These results support the utility of this model for the design of FUS waveforms for neuromodulation.

**Fig. 4.**
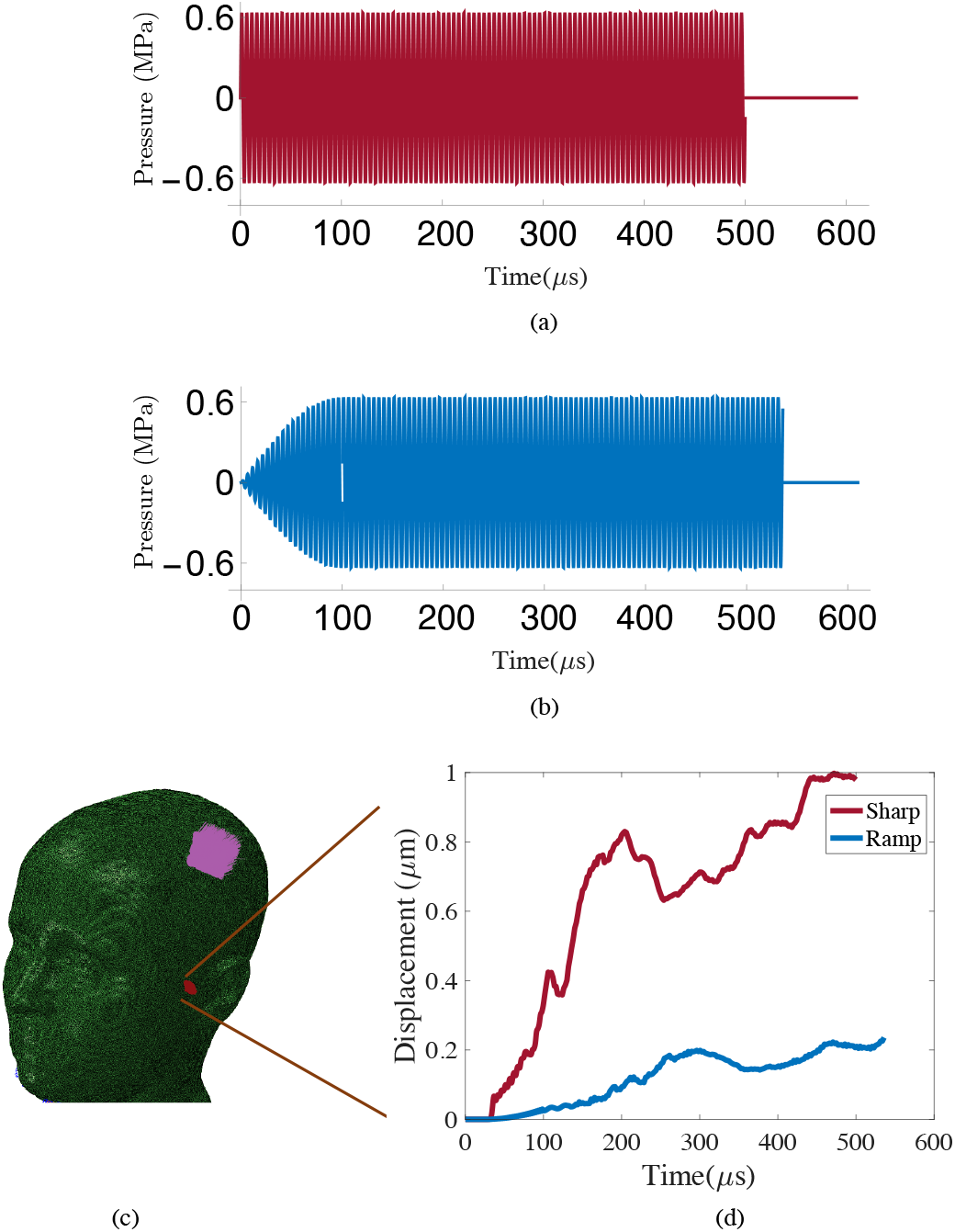
Displacements in a zone in the inner ear resulting from sharp and ramped ultrasound application at 200 kHz to a region of the frontal lobe of human head. (a) The sharp waveform has an amplitude of 0.6 MPa, starting and ending abruptly after 0.5 ms. (b) In the ramped waveform, the maximum 0.6 MPa amplitude is reached gradually over 0.1 ms; it is then continued long enough (0.536 ms) so that both waveforms have the same total energy. (c) Model of the human cranium showing the location of FUS application and the zone of the inner ear where displacements are quantified. (d) Envelope of the family of displacement magnitude curves in the inner ear in response to sharp and ramped pulses.

In summary, our results establish a computational modeling approach incorporating the solid mechanics of cranial tissues in addition to acoustics, and show that this multi-physical combination is essential to fully capture the biophysical effects of transcranial FUS. In the scenario examined in this work, our model demonstrates that the skull can act as a waveguide conducting ultrasound-induced shear waves to the ear, explaining a potential source of auditory side effects in FUS neuromodulation. More generally, the ability of bone to serve as a naturally embedded waveguide for ultrasound-induced shear waves could have implications in multiple other biomedical uses of ultrasound.

## Acknowledgements

The authors thank Dr. Richard Chadwick for helpful discussion. The authors also thank Dr. Hongsun Guo for insightful discussions and input. This project was supported by National Institute of Health grant 1RF1MH117080.

The authors declare no competing financial interest.

## Notes

### Competing Interest Statement

The authors have declared no competing interest.

### Summary of Updates

There was a typo in the abstract, which is fixed in the updated version. Also acknowledgement is added.

## REFERENCES

[1] F. Fry, Ultrasound in medicine & biology 3, 179 (1977).

[2] K. Hynynen and F. A. Jolesz, Ultrasound in medicine & biology 24, 275 (1998).

[3] M. Tanter, J.-L. Thomas, and M. Fink, The Journal of the Acoustical Society of America 103, 2403 (1998).

[4] K. Hynynen, N. McDannold, N. Vykhodtseva, and F. A. Jolesz, Radiology 220, 640 (2001).

[5] S. Mitragotri, Nature Reviews Drug Discovery 4, 255 (2005).

[6] T. Deffieux, J.-L. Gennisson, M. Tanter, M. Fink, and A. Nordez, Applied physics letters 89, 184107 (2006).

[7] Y. Li, H. Zhang, C. Kim, K. H. Wagner, P. Hemmer, and L. V. Wang, Applied physics letters 93, 011111 (2008).

[8] Y.-S. Tung, F. Marquet, T. Teichert, V. Ferrera, and E. E. Konofagou, Applied physics letters 98, 163704 (2011).

[9] J. Ylitalo, J. Koivukangas, and J. Oksman, IEEE Transactions on Biomedical Engineering 37, 1059 (1990).

[10] A. Fenster, D. B. Downey, and H. N. Cardinal, Physics in medicine & biology 46, R67 (2001).

[11] Y. Younan, T. Deffieux, B. Larrat, M. Fink, M. Tanter, and J.-F. Aubry, Medical physics 40, 082902 (2013).

[12] S.-S. Yoo, A. Bystritsky, J.-H. Lee, Y. Zhang, K. Fischer, B.-K. Min, N. J. McDannold, A. Pascual-Leone, and F. A. Jolesz, Neuroimage 56, 1267 (2011).

[13] Y. Tufail, A. Matyushov, N. Baldwin, M. L. Tauchmann, J. Georges, A. Yoshihiro, S. I. H. Tillery, and W. J. Tyler, Neuron 66, 681 (2010).

[14] P. P. Ye, J. R. Brown, and K. B. Pauly, Ultrasound in medicine & biology 42, 1512 (2016).

[15] W. Legon, T. F. Sato, A. Opitz, J. Mueller, A. Barbour, A. Williams, and W. J. Tyler, Nature neuroscience, 17(2) (2014).

[16] R. L. King, J. R. Brown, and K. B. Pauly, Ultrasound in medicine & biology, 40(7) (2014).

[17] R. L. King, J. R. Brown, W. T. Newsome, and K. B. Pauly, Ultrasound in medicine & biology, 39(2) (2013).

[18] W. Lee, H. C. Kim, Y. Jung, I. U. Song, Y. A. Chung, S. S. Yoo, Scientific reports, 5:8743 (2015).

[19] W. Lee, H. C. Kim, Y. Jung, Y. A. Chung, I. U. Song, J. H. Lee, S. S. Yoo, Scientific reports, 6(1) (2016).

[20] N. Wattiez, C. Constans, T. Deffieux, P. M. Daye, M. Tanter, J. F. Aubry, P. Pouget, Brain stimulation, 10(6) (2017).

[21] L. Verhagen, C. Gallea, D. Folloni, C. Constans, D. E. Jensen, H. Ahnine, L. Roumazeilles, M. Santin, B. Ahmed, S. Lehéricy, M. C. Klein-Flügge, Elife, 8, e40541 (2019).

[22] D. Folloni, L. Verhagen, R. B. Mars, E. Fouragnan, C. Constans, J. F. Aubry, M. F. Rushworth, J. Sallet, Neuron, 101(6) (2019).

[23] T. Sato, M. G. Shapiro, and D. Y. Tsao, Neuron 98, 1031 (2018).

[24] H. Guo, M. Hamilton II, S. J. Offutt, C. D. Gloeckner, T. Li, Y. Kim, W. Legon, J. K. Alford, and H. H. Lim, Neuron 98, 1020 (2018).

[25] Braun, V., Blackmore, J., Cleveland, R.O. and Butler, C. R., 2020. Transcranial ultrasound stimulation in humans is associated with an auditory confound that can be effectively masked. bioRxiv.

[26] J. L. Robertson, B. T. Cox, J. Jaros, and B. E. Treeby, The Journal of the Acoustical Society of America 141, 1726 (2017).

[27] A. Jerusalem, Z. Al-Rekabi, H. Chen, A. Ercole, M. Malboubi, M. Tamayo-Elizalde, L. Verhagen, and S. Contera, Acta biomaterialia (2019).

[28] J. Weickenmeier, C. Butler, P. Young, A. Goriely, and E. Kuhl, Computer Methods in Applied Mechanics and Engineering 314, 180 (2017).

[29] Y.-C. Lu, N. P. Daphalapurkar, A. Knutsen, J. Glaister, D. Pham, J. Butman, J. L. Prince, P. Bayly, and K. Ramesh, Annals of biomedical engineering 47, 1923 (2019).

[30] S. Ganpule, N. P. Daphalapurkar, K. T. Ramesh, A. K. Knutsen, D. L. Pham, P. V. Bayly, and J. L. Prince, Journal of neurotrauma 34, 2154 (2017).

[31] R. T. Miller, S. S. Margulies, M. Leoni, M. Nonaka, X. Chen, D. H. Smith, and D. F. Meaney, SAE transac-tions, 2798 (1998).

[32] T. El Sayed, A. Mota, F. Fraternali, and M. Ortiz, Journal of Biomechanics 41, 1458 (2008).

[33] T. El Sayed, A. Mota, F. Fraternali, and M. Ortiz, Computer Methods in Applied Mechanics and Engineering 197, 4692 (2008).

[34] L. A. Mihai, L. Chin, P. A. Janmey, and A. Goriely, Journal of The Royal Society Interface 12, 20150486 (2015).

[35] S. Budday, T. C. Ovaert, G. A. Holzapfel, P. Steinmann, and E. Kuhl, Archives of Computational Methods in Engineering, 1 (2019).

[36] S. Budday, G. Sommer, C. Birkl, C. Langkammer, J. Haybaeck, J. Kohnert, M. Bauer, F. Paulsen, P. Stein-mann, E. Kuhl, et al., Acta biomaterialia 48, 319 (2017).

[37] S. Budday, G. Sommer, G. Holzapfel, P. Steinmann, and E. Kuhl, Journal of the mechanical behavior of biomedi-cal materials 74, 463 (2017).

[38] S. Chatelin, A. Constantinesco, and R. Willinger, Biorheology 47, 255 (2010).

[39] A. P. Sarvazyan, O. V. Rudenko, S. D. Swanson, J. B. Fowlkes, and S. Y. Emelianov, Ultrasound in medicine & biology 24, 1419 (1998).

[40] E. H. Clayton, G. M. Genin, and P. V. Bayly, Journal of The Royal Society Interface 9, 2899 (2012).

[41] J. Achenbach, Wave propagation in elastic solids, Vol. 16 (Elsevier, 2012).

[42] K. Nightingale, S. McAleavey, and G. Trahey, Ultrasound in medicine & biology 29, 1715 (2003).

[43] J. Ophir, I. Cespedes, H. Ponnekanti, Y. Yazdi, and X. Li, Ultrasonic imaging 13, 111 (1991).

[44] J. Bercoff, M. Tanter, and M. Fink, IEEE transactions on ultrasonics, ferroelectrics, and frequency control 51, 396 (2004).

[45] A. Warner, J. Tate, B. Burton, and C. R. Johnson, bioRxiv, 552190 (2019).

[46] H. Mao, L. Zhang, B. Jiang, V. V. Genthikatti, X. Jin, F. Zhu, R. Makwana, A. Gill, G. Jandir, A. Singh, et al., Journal of biomechanical engineering 135, 111002 (2013).

[47] J. D. Stitzel, S. M. Duma, J. M. Cormier, and I. P. Herring, A nonlinear finite element model of the eye with experimental validation for the prediction of globe rupture, Tech. Rep. (SAE Technical Paper, 2002).

[48] J. Newman, C. Barr, M. C. Beusenberg, E. Fournier, N. Shewchenko, E. Welbourne, and C. Withnall, in Pro-ceedings of the International Research Council on the Biomechanics of Injury conference, Vol. 28 (International Research Council on Biomechanics of Injury, 2000).

[49] L. Zhang, K. H. Yang, and A. I. King, J. Biomech. Eng. 126, 226 (2004).

[50] C. Deck and R. Willinger, International Journal of Crashworthiness 13, 667 (2008).

[51] R. M. Wright and K. Ramesh, Biomechanics and modeling in mechanobiology 11, 245 (2012).

[52] T. El Sayed, A. Mota, F. Fraternali, and M. Ortiz, Computer Methods in Applied Mechanics and Engineering 197, 4692 (2008).

[53] B. Dogdas, D. Stout, A. F. Chatziioannou, and R. M. Leahy, Physics in Medicine & Biology 52, 577 (2007).

[54] M. Mohammadjavadi, P. P. Ye, A. Xia, J. Brown, G. Popelka, and K. B. Pauly, Brain stimulation, 12(4), pp.901–910 (2019).

[55] T. Deffieux, Y. Younan, N. Wattiez, M. Tanter, P. Pouget, and J. F. Aubry., Current Biology, 23(23), pp.2430–2433 (2013).

[56] G. Von Békésy, and E. G. Wever. Experiments in hearing. New York: McGraw-Hill (1960).

[57] P. M. Morse, K. U. Ingard. Theoretical acoustics. Princeton university press; 1986.

